# Zika virus-induced TNF-α signaling dysregulates expression of neurologic genes associated with psychiatric disorders

**DOI:** 10.1101/2021.11.15.468744

**Authors:** Po-Lun Kung, Tsui-Wen Chou, Marissa Lindman, Nydia P. Chang, Benjamin D. Buckley, Colm Atkins, Brian P. Daniels

## Abstract

Zika virus (ZIKV) is an emerging flavivirus of global concern. ZIKV infection of the central nervous system has been linked to a variety of clinical syndromes, including microcephaly in fetuses and rare but serious neurologic disease in adults. However, the potential for ZIKV to influence brain physiology and host behavior following recovery from apparently mild or subclinical infection is less well understood. Furthermore, though deficits in cognitive function are well-documented following recovery from neuroinvasive viral infection, the potential impact of ZIKV on other host behavioral domains has not been thoroughly explored. In our study, we performed transcriptomic profiling of primary neuron cultures following ZIKV infection, which revealed altered expression of key genes associated with major psychiatric disorders, such as bipolar disorder and schizophrenia. Gene ontology enrichment analysis also revealed significant changes in gene expression associated with fundamental neurobiological processes, including neuronal development, neurotransmission, and others. These alterations to neurologic gene expression were also observed in the brain *in vivo* using an immunocompetent mouse model of ZIKV infection. Mechanistic studies identified TNF-α signaling via TNFR1 as a major regulatory mechanism controlling ZIKV-induced changes to neurologic gene expression. Our studies reveal that cell-intrinsic innate immune responses to ZIKV infection profoundly shape neuronal transcriptional profiles, highlighting the need to further explore associations between ZIKV infection and disordered host behavioral states.

## Introduction

Zika virus (ZIKV) is a mosquito-borne pathogen of global concern (1). Like many other members of the genus *Flaviviridae*, ZIKV is both neuroinvasive and neurotropic (2). Infection of the central nervous system has been linked to diverse clinical syndromes, including severe congenital neurodevelopmental abnormalities in infants following vertical infection *in utero* (3–5). Severe neurologic disease is less frequent in adults, though cases of encephalitis, myelitis, and, more commonly, peripheral neuropathy have been reported (2, 6). While research on ZIKV pathogenesis to date has heavily focused on severe neurologic disease, it remains unclear whether ZIKV accesses the central nervous system during mild and/or subclinical infection, though data from animal models suggest this is probable (7–9). Even in the case of established neuroinvasive infection, the long-term neurologic consequences that follow viral clearance and recovery remain poorly understood. However, recent evidence suggests that a range of potential neurologic sequalae may occur in the postinfectious brain, including changes to host behavior (10–13).

In particular, recovery from neuroinvasive infection by flaviviruses, including ZIKV, has been associated with neurocognitive deficits (14–17). These effects have been attributed in part to the activities of immune cells, including both T cells and microglia, which act in concert to aberrantly prune neuronal synapses following flavivirus recovery (18–20). Flavivirus infection has also been shown to alter neurodevelopmental programs (21–24), including adult neurogenesis (25–27), a feature of flavivirus infection that contributes to altered learning and memory following recovery in rodent models (28). Cognitive decline, including persistent memory loss, is also a common occurrence in human patients recovering from flavivirus encephalitis (29–31).

Despite these insights into the cognitive consequences of flavivirus infection, the potential impact of these viruses on other behavioral domains remains relatively unexplored. The multifaceted impact of neurotropic flaviviruses on a diverse array of neurologic functions suggests that such infections may also promote or exacerbate neuropsychiatric conditions, including mood and psychotic disorders. Indeed, depression is another common behavioral symptom reported in patients recovering from flavivirus infection (32–34). Case reports have also documented the appearance of psychotic symptoms, including hallucinations, in adult patients infected with ZIKV (35, 36). However, the cellular and molecular mechanisms that underlie these effects remain unknown. In particular, the potential for ZIKV infection to impact host behavior due to cell intrinsic effects on neuronal gene expression has been relatively unexplored.

In this study, we examined how ZIKV infection in neurons impacted the expression of key neurologic genes that promote homeostatic neural function, as well as genes associated with disordered behavioral states. Transcriptomic profiling of primary cortical neurons following ZIKV infection revealed altered expression of a large number of genes associated with psychiatric disorders, including autism, depression, and schizophrenia. Moreover, unbiased gene ontology enrichment analysis revealed that ZIKV infection disproportionately impacted expression of genes associated with neurotransmission and neurodevelopment. These patterns of altered gene expression were also observed *in vivo*, using an established model of CNS ZIKV infection in immunocompetent mice. Observed changes in gene expression were due, at least in part, to innate cytokine signaling via tumor necrosis factor receptor-1 (TNFR1). Our data describe a mechanism linking the cell intrinsic innate immune response to ZIKV with dysregulation of a diverse array of neurologic gene pathways, opening new avenues of inquiry into the effect of flavivirus infection on host behavior.

## Materials and Methods

### Viruses

ZIKV strain MR766 was originally provided by Dr. Andrew Oberst, University of Washington. Viral stocks were generated by infecting Vero cells (MOI 0.01) and harvesting supernatants at 72hpi. Viral titers of stocks were determined via plaque assay on Vero cells (ATCC, #CCL-81). Cells were maintained in DMEM (Corning #10-013-CV) supplemented with 10% Heat Inactivated FBS (Gemini Biosciences #100-106), 1% Penicillin-Streptomycin-Glutamine (Gemini Biosciences #400-110), 1% Amphoteracin B (Gemini Biosciences #400-104), 1% Non Essential Amino Acids (Cytiva #SH30238.01), and 1% HEPES (Cytiva SH30237.01). Plaque assay basal media was 10X EMEM (Lonza # 12-684F) adjusted to 1X and supplemented with 2% Heat Inactivated FBS (Gemini Biosciences #100-106), 1% Penicillin-Streptomycin-Glutamine (Gemini Biosciences #400-110), 1% Amphoteracin B (Gemini Biosciences #400-104), 1% Non Essential Amino Acids (Cytiva #SH30238.01), and 1% HEPES (Cytiva SH30237.01), 0.75% Sodium Bicarbonate (VWR #BDH9280) and 0.5% Methyl Cellulose (VWR #K390). Plaque assays were developed 4dpi by removal of overlay media and staining/fixation using 10% neutral buffered formalin (VWR #89370) and 0.25% crystal violet (VWR #0528).

### Cell culture experiments

Primary cerebral cortical neurons were generated using E15 embryos as described (37). Cells were maintained on cell culture treated multiwell dishes supplemented by coating with 20μg/mL Poly-L-Lysine (Sigma-Aldrich, #9155). Neurobasal Plus + B-27 supplement was used for all experiments (Thermo-Fisher Scientific, #A3582901). All primary mouse cells were generated using pooled tissues derived from both male and female animals. For ZIKV infection experiments, primary neuron cultures were infected at an MOI of 0.1.

### Quantitative real-time PCR

Total RNA from cultured cells was isolated with Qiagen RNeasy mini extraction kit (Qiagen, #74106) following the manufacturer’s protocol. RNA concentration was measured with a Quick Drop device (Molecular Devices). cDNA was subsequently synthesized with qScript cDNA Synthesis Kit (Quantabio, #95048). qPCR was performed with SYBR Green Master Mix (Applied Biosystems, #A25742) using a QuantStudio5 instrument (Applied Biosystems). Cycle threshold (CT) values for analyzed genes were normalized to CT values of the housekeeping gene 18S (CT_Target_ − CT_18S_ = ΔCT). Data were further normalized to baseline control values (ΔCT_experimental_ − ΔCT_control_ = ΔΔCT). Primers were designed using Primer3 (https://bioinfo.ut.ee/primer3/) against murine genomic sequences. A list of primer sequences used in the study appear in **Supplemental Table 1**.

### Murine model of ZIKV infection

C57BL/6J mice were bred in-house for all experiments. All animals were housed under pathogen-free conditions in the animal facilities in Nelson Biological Laboratories at Rutgers University. Both male and female mice were inoculated intracranially (10 μl) with 10^4^ PFU of ZIKV, as described previously (8). ZIKV strain MR-766 was used in all experiments.

### Tissue preparation

All tissues harvested from mice were extracted following extensive cardiac perfusion with 30 mL of sterile PBS. Extracted tissues were weighed and homogenized using 1.0 mm diameter zirconia/silica beads (Biospec Products, #11079110z) in sterile PBS for ELISA (VWR #L0119) or TRI Reagent (Zymo, #R2050-1) for gene expression analysis. Homogenization was performed in an Omni Beadrupter Elite for 2 sequential cycles of 20 seconds at a speed of 4 m/s. Total RNA was extracted using Zymo Direct-zol RNA Miniprep kit, as per manufacturer instructions (Zymo, #R2051).

### ELISA

A TNF-α sandwich ELISA kit (EBioscience, #MTA00B) was used for detection of cytokine levels in cell culture supernatants and brain tissue homogenates. Colorimetric reading of ELISA plates was performed with a microplate reader and Gen5 software (BioTek Instruments, Inc.).

### Neutralizing-antibody studies

Neutralizing-antibody studies were performed after 30 minutes of pretreatment with purified anti-mouse TNFR1 (Invitrogen, # 16-1202-85) and anti-mouse IFNAR1 (Leinco, # I-400) antibodies. IgG isotype antibodies (eBioscience, # eBio299Arm; Leinco, # I-443) were used as controls.

### Curation of psychiatric disorder-associated gene list

Genes associated with autism spectrum disorder (ASD) for our bioinformatics study were identified using the Sfari Gene database (https://gene.sfari.org) (38). The Sfari database includes a ranked list of genes with known associations to ASD. We included genes within the top 3 levels of evidential strength of association (syndromic, category 1, and category 2). All genes within this curation have at least two reported de novo likely-gene-disrupting mutations. We were unable to identify similar database resources for other psychiatric disorders. We thus assembled gene lists for additional disorders by consultation of recent and/or highly cited literature in these areas, including metanalyses and systematic reviews. More information about our gene list can be found in **Supplemental Table 2**.

### Bioinformatics and statistical analysis

Secondary analysis of our previously published microarray dataset (accession # GSE122121) was performed in GEO2R and the GO Enrichment Analysis tool (https://geneontology.org). Biological pathways were defined using the PANTHER (Protein Analysis Through Evolutionary Relationships) classification system. Corrected *p* values (false discovery rate) were determined using the Benjamini & Hochberg procedure. For molecular biology assays, two-way analysis of variance (ANOVA) with Sidak’s correction for multiple comparisons was performed using GraphPad Prism Software v8 (GraphPad Software, San Diego, CA). *P* < 0.05 was considered statistically significant. Data points in all experiments represent biological replicates unless otherwise noted.

## Results

We first assessed whether ZIKV infection in neurons resulted in altered expression of genes associated with abnormal and/or disordered behavioral states. To do so, we first generated a list of 676 genes that have been linked in previous studies to psychiatric disorders, including autism, attention deficit hyperactivity disorder (ADHD), bipolar disorder, major depressive disorder, and schizophrenia (39–86) **(Supplemental Table 2)**. This list of genes includes a combination of known risk genes as well as genes associated with behavioral abnormalities in each of the above human disorders and related animal models. While we stress that the list is not designed to be comprehensive or definitive, it serves as a starting point for probing the behavioral consequences of neuronal ZIKV infection. We assessed the impact of ZIKV infection on the expression of these genes using a dataset previously published by our group and others in which primary cortical neurons derived from C57BL/6J mice were infected with 0.01 MOI ZIKV-MR766 (8). Gene expression in this study was profiled via microarray analysis at 24h following infection.

Cross-comparison of the differentially expressed genes (DEGs) from the microarray analysis with our curated gene list revealed that 181 out of 676 (26.8%) genes exhibited significant differential expression following ZIKV infection **(Figure 1A)**. Of these significant DEGs, 85 (46.96%) were downregulated by ZIKV infection, while 96 (53.04%) were upregulated by ZIKV infection. The 13 genes with the lowest p value in this analysis included *Nr4a2*, *Itgb3*, and *Slc7a5*, which were downregulated by ZIKV infection, along with *Nlgn1*, *Myt1l*, *Dpp6*, *Thsd7a*, *Hepcam*, *Syn3*, *Cacna1e*, *Met*, *Pyhin1*, and *Tor3a*, which were upregulated by ZIKV infection **(Figure 1B)**. The functions of these genes are summarized in **Figure 1C**, and generally include synaptic function, ion channel physiology, and neurodevelopmental processes. None of the specific psychiatric disorder gene lists were significantly overrepresented among the list of significant DEGs **(Figure 1D)**. Together, these data suggest that ZIKV infection in neurons dysregulates expression of a broad set of genes associated with abnormal or pathologic behavioral states.

**Figure 1:**
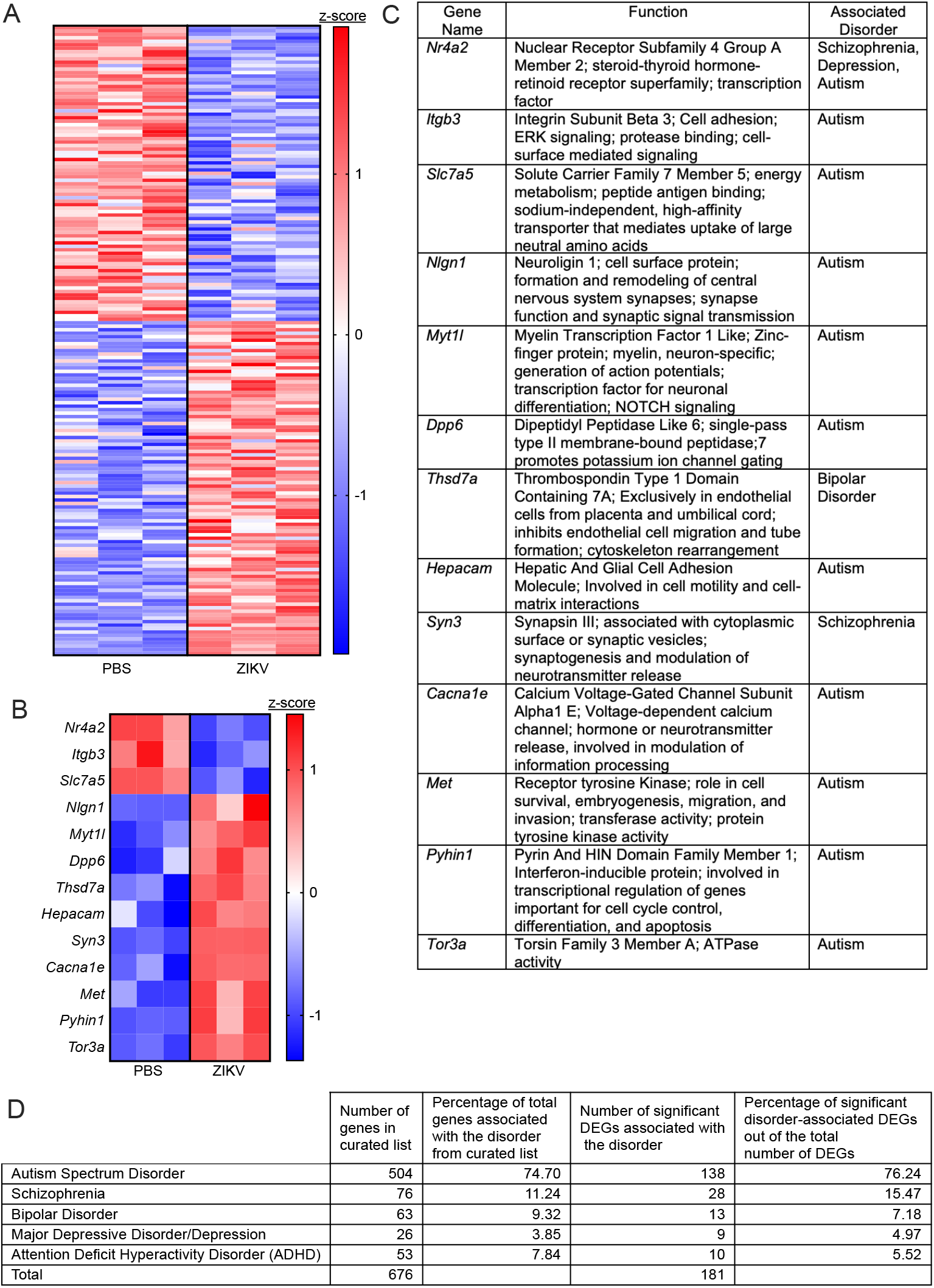
ZIKV infection in neurons dysregulates expression of a broad set of genes associated with abnormal or pathologic behavioral states. **A)** Heatmap depicting relative expression values of 181 candidate genes associated with psychiatric disorders. Values are derived from microarray analysis of primary cortical neurons 24h following ZIKV-MR766 infection (MOI 0.1) or PBS control treatment. **B-C)** Expression values **(B)** of the top 13 significant differentially expressed genes (DEGs) in our microarray analysis (identified by lowest p values). Table in **(C)** lists known functions and associated disorders for these genes. **D)** Descriptive statistics for the curated psychiatric disorder-associated gene list and the significant DEGs observed for each disorder. Data in (**A**) and (**B**) represent normalized and z-transformed expression values. Data in **(A)** include all genes with a False Discovery Rate (FDR) <0.1.

To better understand the consequences of ZIKV infection on neurologic gene expression, we next took an unbiased approach by performing gene ontology (GO) enrichment analysis on the DEGs derived from our microarray dataset. Similar to our results using the curated psychiatric disorder-associated gene list, this unbiased analysis revealed significant enrichment of several GO terms related to neurotransmission, neuronal stress responses, and neurodevelopment, a subset of which are highlighted in **Figure 2A-B**. Notably, regulatory pathways influencing ion transport and ion homeostasis were particularly enriched in our dataset. We next questioned whether there were clear patterns in the direction of differential expression among the significantly enriched GO terms. Heatmaps depicting the expression of genes associated with several representative GO terms are shown in **Figure 2C-E**, each of which reveal a mixed set of both up- and down-regulated genes. These findings suggest that the impact of ZIKV infection on the transcriptional state of neurons likely involves complex alterations to a variety of fundamental neurobiological processes.

**Figure 2:**
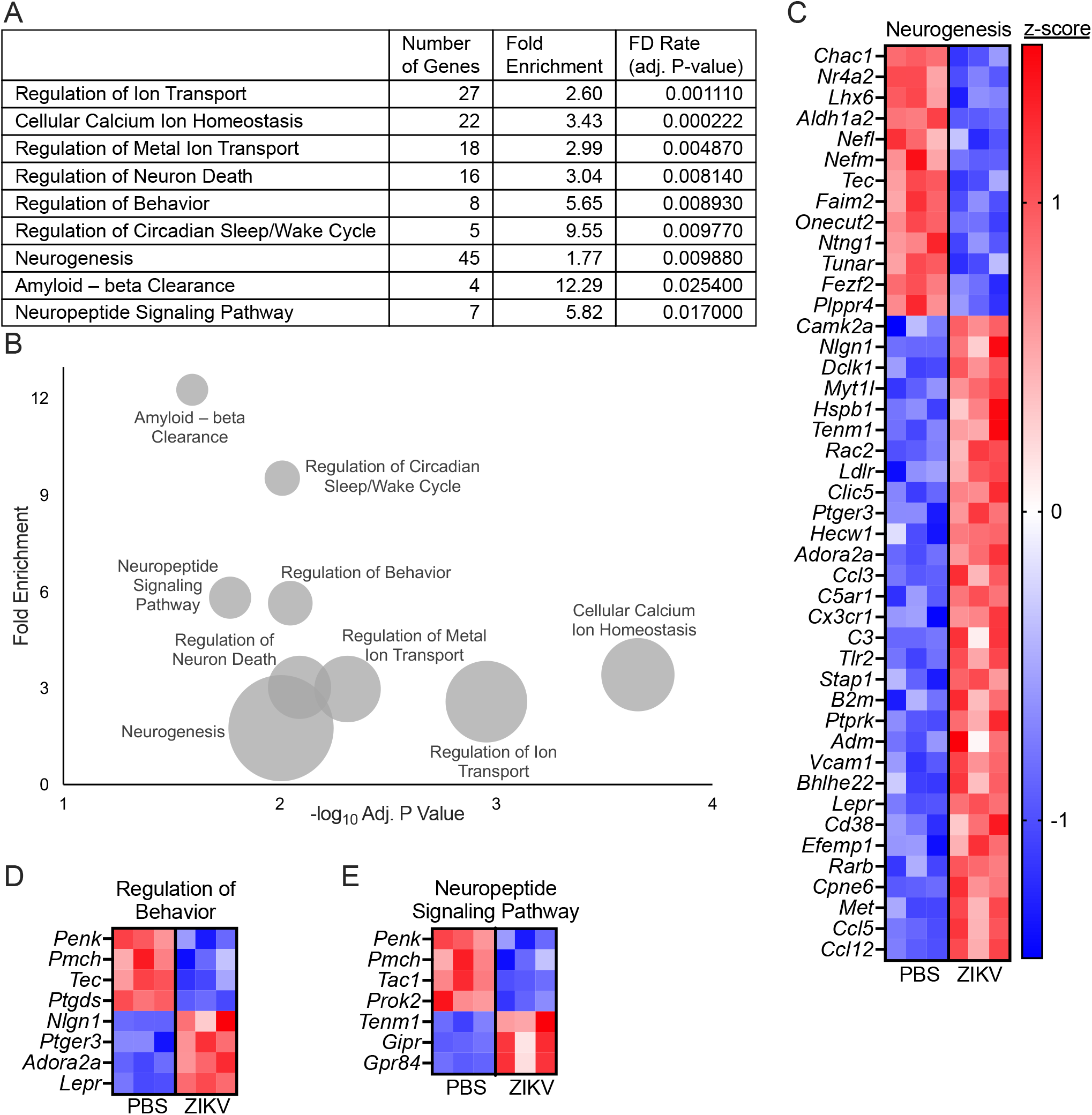
ZIKV infection impacts transcriptional pathways associated with fundamental neurobiological processes. **A-B)** Selected overrepresented GO terms obtained from GO enrichment analysis of DEGs resulting ZIKV-MR766 infection in primary cortical neurons. Tabular results (**A**) are graphically represented in a bubble plot (**B**) to demonstrate associations between fold enrichment, FDR, and number of associated genes for each GO term. **C-E)** Heatmaps depicting expression values for genes associated with neurogenesis (**C**), regulation of behavior (**D**), and neuropeptide signaling pathway (**E**). Data in (**C**), (**D**) and (**E**) represent normalized and z-transformed expression values.

While our microarray data revealed profound alterations to neurologic gene expression in primary neuronal culture following ZIKV infection, we next wanted to assess whether similar changes to gene expression occur *in vivo.* To do so, we inoculated male and female wildtype (C57BL/6J) mice intracranially with 10^4^ pfu ZIKV-MR766. We performed these studies in separate cohorts of adolescent (3 week old) and adult (8 week old) animals to account for potential differences in the expression of neurologic genes across development **(Figure 3A)**. On days 2 and 4 following infection, we harvested brains and used qRT-PCR to assess the expression of a panel of genes derived from the top DEGs identified in our microarray analysis. This candidate gene panel included genes from both our curated psychiatric disorder-associated gene list, as well as DEGs from the highly enriched GO terms identified in Figure 2.

**Figure 3:**
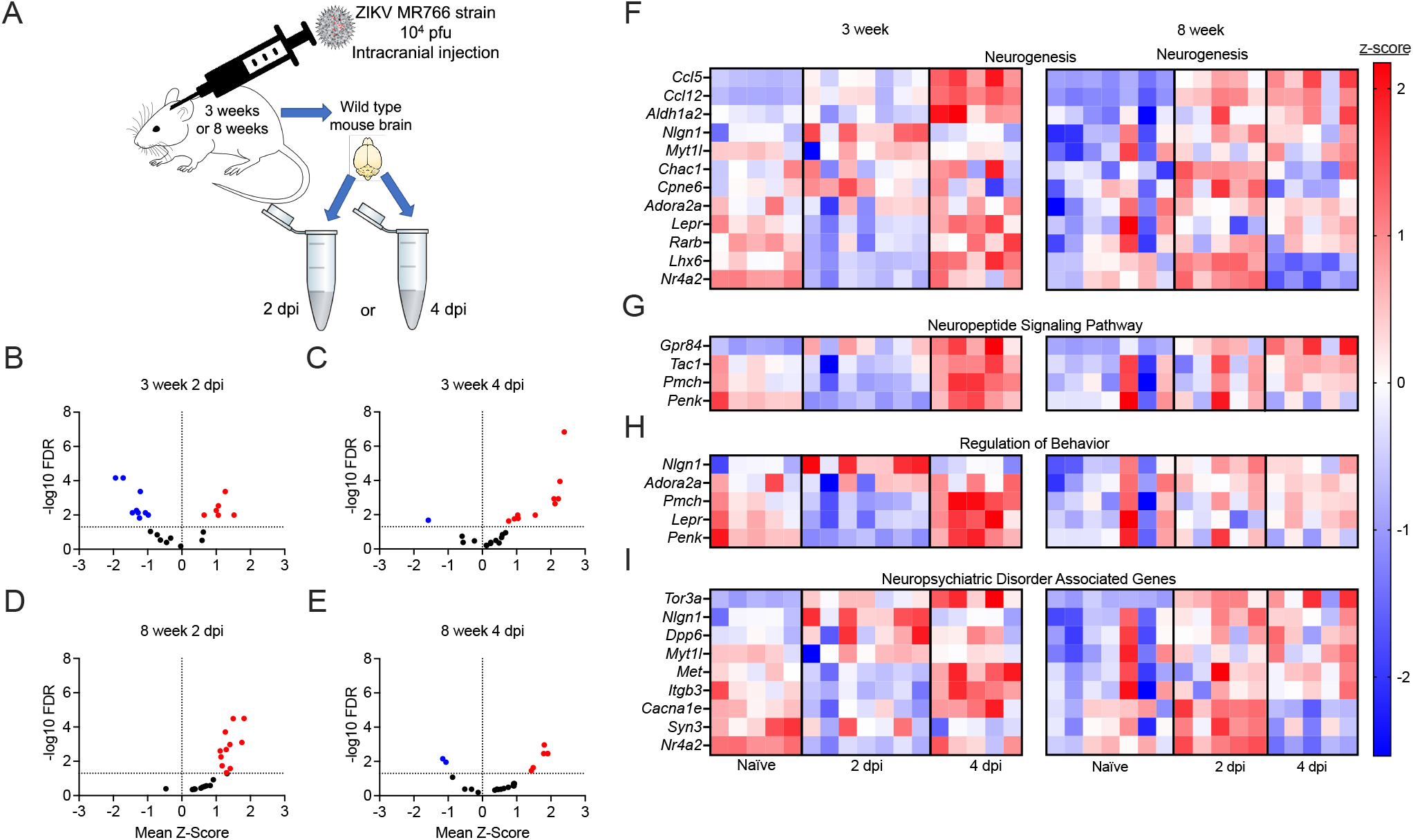
Gene expression associated with disease-relevant neurologic pathways is significantly dysregulated following ZIKV infection of the brain *in vivo*. **A)** Schematic of experimental design of *in vivo* murine model. C57/BL/6J WT mice were infected with 10^4^ pfu ZIKV-MR766 via intracranial inoculation at 3 and 8 weeks of age. Whole brain homogenates were collected at 2 or 4 days post infection (dpi). **B-E)** Expression profiles of 23 candidate genes assessed by qRT-PCR analysis are described and separated into 3-week mice at 2dpi (**B**), 3-week mice at 4dpi (**C**), 8-week mice at 2dpi (**D**), and 8-week mice at 4dpi (**E**). Significant differences (*p* < 0.05) are noted in red (upregulated DEGs) or blue (downregulated DEGs). **F-I)** Heatmaps showing individual expression values per mouse for candidate genes from microarray and GO enrichment analyses in 3 week old adolescent and 8 week old adult mouse brains following ZIKV-MR766 infection. Relative gene expression values are reported for naïve, 2 dpi, or 4 dpi groups. Data in (**F**), (**G**), (**H**) and (**I**) represent normalized and z-transformed values of qRT-PCR expression data. n= 5-7 mice/group.

These experiments revealed that a majority of genes in our panel did exhibit differential expression at the whole-brain level following intracranial ZIKV infection *in vivo*. However, the magnitude and direction of differential expression exhibited complex patterns that differed across time post infection and between 3 week old and 8 week old animals. In particular, 3 week old animals exhibited a mix of significantly downregulated and upregulated expression of selected neurologic genes 2 days post infection (dpi), with a marked shift to primarily upregulated expression at 4dpi **(Figure 3B-C)**. In contrast, 8 week old animals exhibited an essentially inverse pattern, with nearly uniform upregulation of significant DEGs at 2dpi, but a mixture of down- and up-regulated DEGs at 4dpi **(Figure 3D-E)**. These patterns of differential expression were evident across GO terms, including neurogenesis **(Figure 3F)**, neuropeptide signaling pathway **(Figure 3G)**, and regulation of behavior **(Figure 3H)**, as well as genes taken from our curated list of psychiatric disorder-associated genes **(Figure 3I)**. Together, these data confirm that gene expression associated with disease-relevant neurologic pathways is significantly dysregulated following ZIKV infection of the brain *in vivo*, but the mechanisms that control these transcriptional responses are under complex regulation by factors that vary with host age.

We next questioned whether innate immune activation following ZIKV infection in neurons may be linked to observed changes in neurologic gene expression. To answer this, we returned to our gene ontology enrichment analysis to identify innate immune signaling pathways that were most significantly impacted by neuronal ZIKV infection. Well-known antiviral cytokine responses were among the most enriched pathways in this analysis, particularly those related to innate cytokines, including type I interferon (IFN), interleukin (IL)-1 and IL-6, and tumor necrosis factor (TNF)-α **(Figure 4A-B)**. Pathways related to signaling by the inflammatory transcription factor nuclear factor kappa B (NF-κB) were also particularly enriched within the list of significant DEGs. We next confirmed that each of these major cytokine responses was induced in the brains of 8 week old mice following intracranial infection with ZIKV. Expression analysis via qRT-PCR showed that ZIKV infection induced significant upregulation of each of the cytokines analyzed, including *Ifna6*, *Ifnb*, *Ifng*, *Il1b*, *Il6*, and *Tnfa* **(Figure 4C-H).** Together, these data confirm that wildtype neurons mount a robust innate immune cytokine response to ZIKV infection, including several cytokines previously established to influence brain function and behavior.

**Figure 4:**
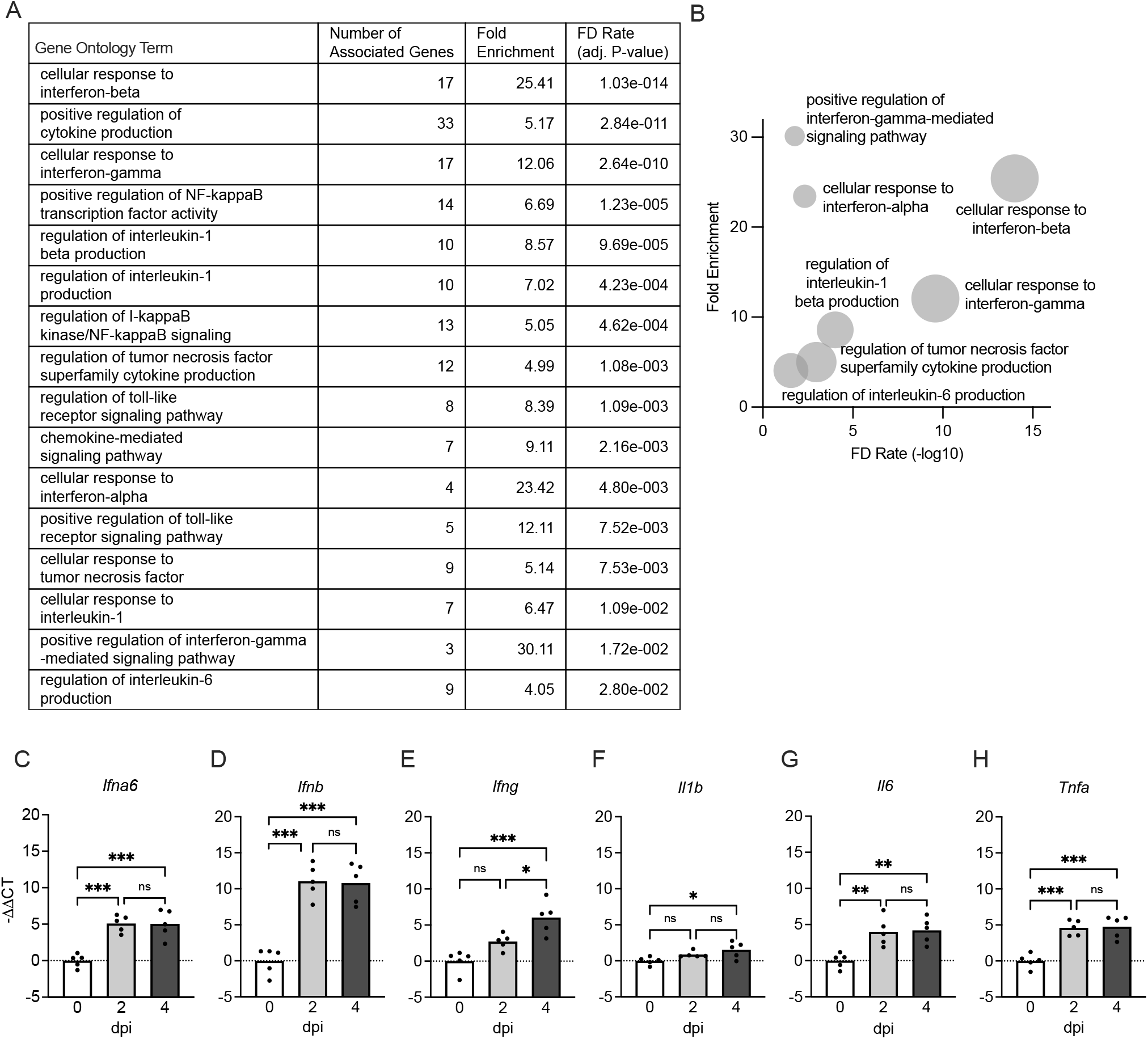
Neurons mount a robust innate immune cytokine response to ZIKV infection. **A-B)** Results of GO enrichmsent analysis of microarray data derived from primary cortical neurons following ZIKV-MR766 infection (MOI 0.1) compared to PBS-treated controls after 24 hours. Selected GO terms focus on cytokine activation or inflammatory transcription factor responses. Tabular results (**A**) are graphically represented in a bubble plot (**B**) to demonstrate associations between fold enrichment, FDR, and number of associated genes for each GO term. **C-H)** qRT-PCR analysis was performed measuring cytokine genes *Ifna6* (**C**), *Ifnb* (**D**), *Ifng* (**E**), *Il1b* (**F**), *Il6* (**G**), *and Tnfa* (**H**) at indicated time points in 8 week old adult mouse brains following intracranial ZIKV-MR766 infection. n=5. ns not significant, * *p* < 0.05, ** *p* < 0.01, *** *p* < 0.001. Bars represent group means.

To examine whether neuronal cytokines may be implicated in ZIKV-induced changes in neurologic gene expression, we generated primary cultures of cortical neurons and examined expression of targets in our candidate gene panel following treatment with exogenous cytokines, including IFN-β, IL-6, and TNF-α. We then compared the direction of differential expression for each gene to that induced by ZIKV to see which, if any, cytokine most closely phenocopied the pattern of gene expression induced by ZIKV infection. While IFN-β and IL-6 did significantly alter the expression of some target genes, significant DEGs following these treatments did not closely follow the pattern of downregulation **(Figure 5A)** or upregulation **(Figure 5B)** induced by ZIKV infection. In contrast, TNF-α treatment induced a strikingly similar pattern of differential expression to that induced by ZIKV infection. Moreover, while IFN-β and IL-6 only induced significant changes in expression of a handful of neurologic genes in our analysis **(Figure 5C-D)**, TNF-α significantly altered 17 out of 18 genes in our panel **(Figure 5E)**, and of these, all but 2 matched the pattern of up- or downregulated expression observed in ZIKV-infected neuronal cultures. These data identified TNF-α signaling as a promising candidate mechanism for the altered neurologic gene expression observed in the setting of neuronal ZIKV infection.

**Figure 5:**
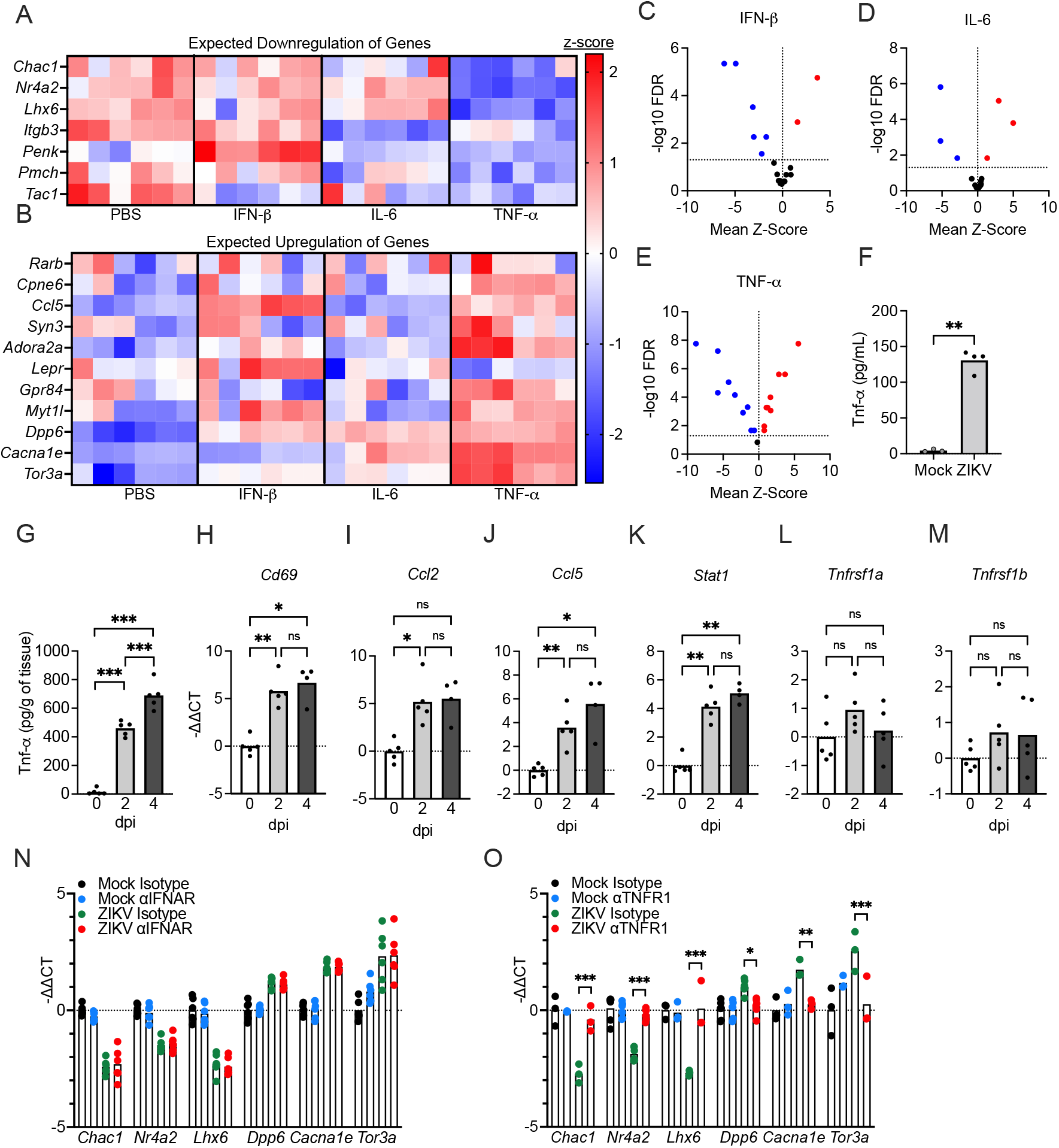
Induction of TNF-α following neuronal ZIKV infection is a major regulatory mechanism that alters expression of genes relevant to neuronal function. **A-B)** Expression values of candidate genes previously shown to be either downregulated (**A**) or upregulated (**B**) by ZIKV infection were measured via qRT-PCR in neuronal cultures treated for 24h with exogenous cytokines: IFN-β, IL-6, or TNF-α. n=6. **C-E)** Volcano plots depicting mean z-score and −log10FDR of representative DEGs in neurons treated with IFN-β(**C**), IL-6 (**D**), or TNF-α(**E**). Significant differences (*p* < 0.05) are noted in red (upregulated DEGs) or blue (downregulated DEGs). n=6. **F-G)** Concentrations of TNF-α in supernatants of *in vitro* neuronal cell cultures infected with ZIKV for 24h (n=4) (**F**) and brains harvested following *in vivo* intracranial infection (n=5) (**G**). Cytokine concentrations were quantified via ELISA assay. **H-M)** qRT-PCR analysis was performed for known TNF-α transcriptional target genes, *Cd69* (**H**), *Ccl2* (**I**), *Ccl5* (**J**), *Stat1* (**K**), and TNF-α receptor genes, *Tnfrsf1a* (**L**) and *Tnfrsf1b* (**M**), in 8 week old adult mouse brains following ZIKV-MR766 infection at 2 or 4 dpi. n=5. **N-O)** qRT-PCR analysis of representative DEGs associated with neurological functions (*Chac1, Nr4a2, Lhx6, Dpp6, Cacna1e, and Tor3a*) following pretreatment with neutralizing antibodies against IFN α/β receptor (IFNAR) (**N**) or TNFR1 (**O**) and subsequent 24h infection with ZIKV-MR766. n=3-6 biological replicates per group. Data in (**A**) and (**B**) represent normalized and z-transformed values of qRT-PCR expression data. ns not significant, * *p* < 0.05, ** *p* < 0.01, *** *p* < 0.001. Bars represent group means.

While we previously confirmed that *Tnfa* was induced at the transcriptional level in the brain *in vivo* following ZIKV infection, we next wanted to confirm that a robust TNF-α-dependent signature could indeed be observed following infection. We thus performed enzyme-linked immunosorbent assay (ELISA) to confirm that TNF-α was upregulated at the protein level in both supernatants of primary neuronal cultures at 24h following infection **(Figure 5F)** and whole brain homogenates derived from 8 week old animals on days 2 and 4 following intracranial ZIKV inoculation **(Figure 5G)**. We also confirmed upregulation of known TNF-α transcriptional targets, including *Cd69*, *Ccl2*, *Ccl5*, and *Stat1*, in the brains of infected 8 week old mice **(Figure 5H-K)**. In contrast, transcript expression of the TNF-α receptors TNFR1 (*Tnfrsf1a*) and TNFR2 (*Tnfrsf1b*) were not altered in the brain following infection **(Figure 5L-M)**, suggesting that enhanced TNF-α signaling in this setting is mediated primarily through induction of cytokine expression. Together, these data confirm that TNF-α signaling is active in both cultured neurons and the brain *in vivo* following ZIKV infection.

To more carefully assess whether TNF-α signaling was required for ZIKV-mediated alterations to neurologic gene expression, we cultured primary cortical neurons and pretreated with neutralizing antibodies against cytokine receptors for 2h prior to infection. After 24h, we then performed qRT-PCR analysis of major DEGs from our microarray analysis to assess the impact of cytokine signaling on ZIKV-induced gene expression. These experiments revealed that blockade of type I IFN signaling via neutralization of the IFN α/β receptor (IFNAR) had no impact on ZIKV-induced changes in expression of the neurologic genes we analyzed **(Figure 5N)**, findings which mirrored our previous result showing that exogenous IFN-β treatment did not phenocopy ZIKV-induced patterns of expression in our target gene list **(Figure 5A-B)**. In contrast, blockade of TNFR1 rescued ZIKV-induced changes in each of the 6 genes we analyzed, including *Chac1*, *Nr4a2*, *Lhx6*, *Dpp6*, *Cacna1e*, and *Tor3a* **(Figure 5O)**. Taken together, these data suggest that the induction of TNF-α following neuronal ZIKV infection is a major regulatory mechanism that alters expression of genes relevant to neuronal function.

## Discussion

Emerging flaviviruses represent a significant and growing challenge to global public health. While most famously associated with rare but severe clinical manifestations, including encephalitis, congenital abnormalities, etc., the consequences of apparently mild and/or asymptomatic infection by neuroinvasive flaviviruses remain poorly understood (2, 87). The observation of behavioral sequalae following recovery from severe flavivirus infections raises the possibility that subclinical neuroinvasive infection may also impact brain function in ways that promote or exacerbate psychiatric disorders. This idea is supported by some case reports (15, 35, 88–90), though, to our knowledge, this hypothesis has not been rigorously tested in the clinical literature. The prevalence of psychiatric sequelae following ZIKV infection, in particular, may be hard to discern due to the relatively low neurovirulence of ZIKV compared to other flaviviruses, resulting in symptoms that may not be severe enough to warrant clinical attention and the documentation of infection status. Our study highlights the need for increased attention to behavioral symptoms in patients who are seropositive for ZIKV and other neuroinvasive flaviviruses, as well as further mechanistic investigation into the cellular and molecular impacts of flavivirus infection on brain physiology and function.

In our study, we show that neurons mount a robust innate cytokine response to ZIKV infection, including a number of cytokines with previously established effects on behavior. A large body of evidence has established that neuroinflammation and inflammatory cytokine signaling is associated with psychiatric disorders, including major depressive disorder (91–94) and schizophrenia (95–97). TNF-α is a major pleiotropic cytokine induced strongly in the CNS by ZIKV and other flaviviruses (98–101). Notably, recent work has described complex neuromodulatory effects of TNF-α signaling, including direct effects on glutamatergic neurotransmission (102–104), neuronal differentiation (105–107), and other fundamental neurologic processes (108–110). In our study, ZIKV-mediated changes to neurologic gene expression greatly overlapped those induced by TNF-α, and gene expression changes induced by ZIKV could be rescued in part by blockade of TNFR1 signaling. These data identify TNF-α as a candidate for further mechanistic investigation of the potential impacts of flavivirus infection on neuronal function.

To date, the most well-described behavioral outcomes of neuroinvasive flavivirus infection in animal models are changes to learning and memory (18, 19, 27, 111–113). While it is clear that a variety of pathogenic processes related both to viral infection and neuroinflammation can impact cognition, comparatively less attention has been devoted to how flavivirus infection impacts other behavioral domains, including mood, affect, and emotional regulation. This discrepancy is likely due, in part, to technical limitations, including difficulty modeling these behavioral domains in rodents and containment issues related to using ABSL2 and ABSL3 pathogens within behavioral laboratories. Nevertheless, our data identify a need for more robust assessment of behavioral changes in models of flavivirus infection, particularly measures of anxiety, fear/avoidance, and other paradigms with relevance to human psychiatric disorders.

Finally, our data add to a growing body of evidence suggesting that the impact of flavivirus infection varies across the lifespan. In our study, intracranial ZIKV infection resulted in very different impacts on neurologic gene expression in adolescent compared to adult animals, suggesting that factors such as developmental states and immune system maturation may significantly influence the neurologic outcomes of flavivirus infection. Recent work has shown that neuroimmune responses to other flaviviruses are impacted by aging (114, 115), and thus differential engagement of cytokine signaling, adaptive immune priming, and blood-brain barrier function may all be relevant variables in determining how flavivirus infection might impact behavior differentially across life stages. Moreover, while the potential for ZIKV to induce severe congenital abnormalities following vertical transmission *in utero* has now been well established, it remains less clear what the impact of ZIKV infection is on apparently developmentally normal fetuses, including those who are exposed to ZIKV late in gestation, when rates of microcephaly and severe birth defects are exceedingly rare (113, 116, 117). Further work will be needed to assess whether ZIKV infection in this context may result in changes to neurodevelopment and brain function that impact behavior postnatally and beyond.

## Declarations

### Ethics Approval and Consent to Participate

All animal experiments were performed with approval of the Rutgers University Institutional Animal Care and Use Committee (IACUC).

### Consent for Publication

Not applicable

### Availability of Data and Materials

All data are available upon reasonable request to the corresponding author. Microarray data used in this study are deposited in NCBI’s Gene Expression Omnibus and can be accessed under accession number GSE122121.

### Contributions

Conceptualization: PLK and BPD; Investigation and analysis: PLK, TWC, ML, NPC, BDB, CA, and BPD.; Writing-original draft: PLK and BPD; Writing-review and editing: PLK, TWC, NPC, CA, and BPD; Supervision: CA and BPD; Funding acquisition: BPD.

## Acknowledgements

Not applicable.

