## Supplemental Material for "Zika virus-induced TNF-α signaling dysregulates expression of neurologic genes associated with psychiatric disorders"

Supplemental Table 1
**Primer Sequences for qRT-PCR**

| **Gene** | **Direction** | **Sequence** |
|---|---|---|
| *Adora2a* | Forward | CACGCAGAGTTCCATCTTCAGC |
| *Adora2a* | Reverse | CCCAGCAAATCGCAATGATGCC |
| *Aldh1a* | Forward | CACAAGACACGAGCCCATTGGA |
| *Aldh1a* | Reverse | GGTTTGATGACCACGGTGTTACC |
| *Cacna1e* | Forward | ATGGACAAGGCTACCACGGAGA |
| *Cacna1e* | Reverse | GACTGGCTTCTCCATCCGTCTT |
| *Ccl12* | Forward | GCTACAGGAGAATCACAAGCAGC |
| *Ccl12* | Reverse | ACGTCTTATCCAAGTGGTTTATGG |
| *Ccl2* | Forward | TGGCTCAGCCAGATGCAGT |
| *Ccl2* | Reverse | TTGGGATCATCTTGCTGGTG |
| *Ccl5* | Forward | CCTGCTGCTTTGCCTACCTCTC |
| *Ccl5* | Reverse | ACACACTTGGCGGTTCCTTCGA |
| *Cd69* | Forward | TTTGGAGGGGTTTCAGTGGT |
| *Cd69* | Reverse | GCTGTCTACACGGAGGAAGT |
| *Chac1* | Forward | TGACCCTCCTTGAAGACCGTGA |
| *Chac1* | Reverse | AGTGTCATAGCCACCAAGCACG |
| *Cpne6* | Forward | TCAGCTTCACGGTGGCTATCGA |
| *Cpne6* | Reverse | CTGTCATAGTCCTGGCAGATACC |
| *Dpp6* | Forward | CAATCCTGACCCTCTGTGATGC |
| *Dpp6* | Reverse | GCCATCTTTGGAGAACACAGGC |
| *Gpr84* | Forward | AAGCCTTCCAGAAGTGCATCGC |
| *Gpr84* | Reverse | CAGAGGAACACTGCGAAGCACA |
| *Icam1* | Forward | AAACCAGACCCTGGAACTGCAC |
| *Icam1* | Reverse | GCCTGGCATTTCAGAGTCTGCT |
| *Ifna6* | Forward | CTTCCACAGGATCACTGTGTACCT |
| *Ifna6* | Reverse | TTCTGCTCTGACCACCTCCC |
| *Ifnb* | Forward | CTGGAGCAGCTGAATGGAAAG |
| *Ifnb* | Reverse | CTT CTC CGT CAT CTC CAT AGG G |
| *Ifng* | Forward | AACGCTACACACTGCATCTTGG |
| *Ifng* | Reverse | GCCGTGGCAGTAACAGCC |
| *Il1b* | Forward | ACCTGTCCTGTGTAATGAAAGACG |
| *Il1b* | Reverse | TGGGTATTGCTTGGGATCCA |
| *Il6* | Forward | CAGAATTGCCATCGTACAACTCTTTTCTCA |
| *Il6* | Reverse | AAGGTCATCATCGTTGTTCATACA |
| *Itgb3* | Forward | GTGAGTGCGATGACTTCTCCTG |
| *Itgb3* | Reverse | CAGGTGTCAGTGCGTGTAGTAC |
| *Lepr* | Forward | CTTTCCTGTGGACAGAACCAGC |
| *Lepr* | Reverse | AGCACTGAGTGACTCCACAGCA |
| *Lhx6* | Forward | AGAGAAAGCACCTCCCAGAA |
| *Lhx6* | Reverse | TGGAGTTCTGAACCAAACCA |
| *Met* | Forward | GTTCTGCTTGGCAACGAGAGCT |

| | | |
|---|---|---|
| *Met* | Reverse | GGAGAATGCACTGTATTGCGTCG |
| *Myt1l* | Forward | TGTCTGGATGCCCGCACAAAGA |
| *Mytl1* | Reverse | CTTCTGTGCGAGTTCCTGTTGC |
| *Nlgn1* | Forward | TGATGGGAGTGTCTTGGCAAGC |
| *Nlgn1* | Reverse | CCGTAGTTTCCTTTGGCAGCCT |
| *Nr4a2* | Forward | GCATACAGGTCCAACCCAGT |
| *Nr4a2* | Reverse | AATGCAGGAGAAGGCAGAAA |
| *Penk* | Forward | TGCAGCCAGGACTGCGCTAAAT |
| *Penk* | Reverse | GATCCTTGCAGGTCTCCCAGAT |
| *Pmch* | Forward | CTCTGGAACAATACAAAAACGACG |
| *Pmch* | Reverse | GGTTTTACAGCCAGACTCAGTGG |
| *Parb* | Forward | GCTTCGTTTGCCAGGACAAGTC |
| *Parb* | Reverse | TGGCATCGGTTCCTAGTGACCT |
| *Stat1* | Forward | GAACGCGCTCTGCTCAA |
| *Stat1* | Reverse | TGCGAATAATATCTGGGAAAGTAA |
| *Syn3* | Forward | CCATCTCTGGAAACTGGAAGGC |
| *Syn3* | Reverse | CGCCGAACATTTCTGAGCAGCT |
| *Tac1* | Forward | TAATGGGCAAGCGGGATGCTGA |
| *Tac1* | Reverse | CCATTAGTCCAACAAAGGAATCTG |
| *Tnfa* | Forward | AAAATTCGAGTGACAAGCCTGTAGC |
| *Tnfa* | Reverse | GTGGGTGAGGAGCACGTAG |
| *Tnfsrsf1a* | Forward | ACGAATCACTCTGCTCCGTG |
| *Tnfsrsf1a* | Reverse | TCGCAAGGTCTGCATTGTCA |
| *Tnfsrsf1b* | Forward | GAGATGCCAAGGTGCCTCAT |
| *Tnfsrsf1b* | Reverse | AACTGGGTGCTGTGGTCAAC |
| *Tor3a* | Forward | GGTGCCTATGTCTCAGCCTCTT |
| *Tor3a* | Reverse | GCCATAGAACTGATGAGAGGGAG |

Supplemental Table 2
**Curated Psychiatric Disorder-Associated Gene List**

| Schizo-phrenia | Bipolar Disorder | Major Depressive Disorder | ADHD | Autism Spectrum Disorder | Implicated gene | Reference(s) |
|---|---|---|---|---|---|---|
| | | | X | | *5HT1B* | (56) |
| | | X | | | *5HTT* | (65) |
| | | | | X | *ACHE* | (39) |
| | | | | X | *ACTB* | (70) |
| | | | | X | *ACTL6B* | (39) |
| | | | | X | *ACY1* | (39) |
| | | | | X | *ADA* | (39) |
| | X | | | | *ADCY2* | (71) |
| | | | | X | *ADCY3* | (39) |
| | | | | X | *ADNP* | (39) |
| | | | X | | *ADRA1A* | (57) |
| | | | X | | *ADRA1B* | (57) |
| | | | | X | *ADSL* | (39) |
| | | | | X | *AFF2* | (39) |
| | | | | X | *AGAP2* | (39) |
| | | | | X | *AGO1* | (39) |
| | | | | X | *AGO2* | (39) |
| | | | | X | *AGO4* | (39) |
| | | | | X | *AHDC1* | (39) |
| | | | | X | *AHI1* | (39) |
| | | | | X | *AHNAK* | (39) |
| | | | | X | *AKAP9* | (39) |
| | | | | X | *ALDH1A3* | (39) |
| | | | | X | *ALDH5A1* | (39) |
| X | | | | | *ALDOA* | (79) |
| | | | | X | *ALG6* | (39) |
| X | | | | | *ALPK3* | (79) |
| | | | | X | *AMPD1* | (39) |
| | X | | | | *AMPD3* | (59) |
| | | | | X | *AMT* | (39) |
| | | | | X | *ANK2* | (39) |
| | | | | X | *ANK3* | (39) |
| | | | X | X | *ANKRD11* | (39) |
| | | | | X | *ANXA1* | (39) |
| | | | | X | *AP1S2* | (39) |

| | | | | | | |
|---|---|---|---|---|---|---|
| | | | | X | *AP2S1* | (39) |
| | | | | X | *APBB1* | (39) |
| | | | | X | *APH1A* | (39) |
| | | X | | | *APOE* | (65) |
| | | | | X | *ARHGEF9* | (39) |
| | | | | X | *ARID1B* | (39) |
| | | | X | | *ARTN* | (53) |
| X | | | | | *ARVCF* | (64) |
| | | | | X | *ARX* | (39) |
| | | | | X | *ASAP2* | (39) |
| | | | | X | *ASH1L* | (39) |
| | | | | X | *ASPM* | (39) |
| | | | | X | *ASTN2* | (39) |
| | | | | X | *ASXL3* | (39) |
| X | | | | | *ATK3* | (79) |
| | | | X | | *ATM* | (57) |
| | | | | X | *ATP10A* | (39) |
| X | | | | | *ATP2A2* | (79) |
| | | | | X | *ATP2B2* | (39) |
| | | | X | | *ATP6V0B* | (53) |
| | | | | X | *ATRX* | (39) |
| | X | | | | *ATXN7L3* | (81) |
| | | | | X | *AUTS2* | (39) |
| | | | | X | *AVPR1A* | (39) |
| | | | | X | *B4GALT2* | (53) |
| | | | | X | *BAZ2B* | (39) |
| | | | | X | *BCKDK* | (39) |
| | | | | X | *BCL11A* | (39) |
| | | | | X | *BCORL1* | (39) |
| | X | X | | | *BDNF* | (65) |
| | | | | X | *BICRA* | (39) |
| X | | | | | *BNIP3L* | (79) |
| | | | | X | *BRAF* | (39) |
| | | | | X | *BRSK2* | (39) |
| | | | | X | *BRWD3* | (39) |
| | | | | X | *BTAF1* | (39) |
| X | | | | | *BTG1* | (79) |
| | | | | X | *C12orf57* | (39) |
| | | | | X | *CACNA1A* | (39) |

| | | | | | | |
|---|---|---|---|---|---|---|
| X | X | | | X | *CACNA1C* | (39), (46), (47) |
| | | | | X | *CACNA1D* | (39) |
| | | | | X | *CACNA1E* | (39) |
| | | | | X | *CACNA1H* | (39) |
| | | | | X | *CACNA2D3* | (39) |
| | X | | | X | *Cacna2d4* | (79) |
| X | | | | X | *CACNB2* | (39) |
| | | | | X | *CAMK2B* | (39) |
| | | | X | | *CAMK2D* | (57) |
| | | | X | | *CAMK2G* | (57) |
| | | | | X | *CAPRIN1* | (39) |
| | | | | X | *CASK* | (39) |
| | | | | X | *CASZ1* | (39) |
| | | X | | | *CATSPER1* | (81) |
| | | | | X | *CC2D1A* | (39) |
| | | | X | | *CCDC24* | (53) |
| | | | | X | *CCNG1* | (39) |
| | | | | X | *CCNK* | (39) |
| | | | | X | *CCT4* | (39) |
| | | | | X | *CDC42BPB* | (39) |
| | | | | X | *CDH13* | (39) |
| | | | | X | *Cdh8* | (74) |
| | | | | X | *CDK13* | (39) |
| X | | | | | *CDK2AP1* | (79) |
| | | | | X | *CDK8* | (39) |
| | | | | X | *CDKL5* | (39) |
| | | | | X | *CELF4* | (39) |
| | | | | X | *CEP135* | (39) |
| | | | | X | *CEP41* | (39) |
| | | | | X | *CGNL1* | (39) |
| | | | | X | *CHAMP1* | (39) |
| | | | | X | *CHD2* | (39) |
| | | | | X | *CHD3* | (39) |
| | | | | X | *CHD7* | (39) |
| | | | | X | *CHD8* | (39) |
| | | | | X | *CHKB* | (39) |
| | | | | X | *CHMP1A* | (39) |
| | | | | X | *CHRNA7* | (39) |
| | | | | X | *CIB2* | (39) |

| | | | | | | |
|---|---|---|---|---|---|---|
| | | | | X | *CIC* | (39) |
| | | | | X | *CLASP1* | (39) |
| X | | | | | *CLCN3* | (51) |
| X | | | | | *CLU* | (79) |
| | | | | X | *CNKSR2* | (39) |
| | | | | X | *CNOT1* | (39), (40) |
| | | | | X | *CNOT3* | (39), (51) |
| | | | X | X | *CNR1* | (39) |
| | | | | X | *CNTN3* | (48) |
| X | | | | X | *CNTN4* | (39) |
| | | | | X | *CNTN5* | (39) |
| | | | | X | *CNTN6* | (39) |
| | | | | X | *CNTNAP2* | (39) |
| | | | | X | *CNTNAP3* | (39) |
| | | | | X | *CNTNAP4* | (39) |
| | | X | X | | *COMT* | (80) |
| | | | | X | *CORO1A* | (39) |
| X | | | | | *CPEB1* | (79) |
| | | | | X | *CPEB4* | (39) |
| X | | | | | *CPT1C* | (79) |
| X | | | | | *CREB3L1* | (79) |
| | | | | X | *CREBBP* | (39) |
| | | | | X | *CSDE1* | (39) |
| | | | | X | *CSNK2A1* | (39) |
| | X | | | | *CSRNP3* | (81) |
| | | | | X | *CTCF* | (39) |
| | | | | X | *CTNNA2* | (39) |
| | | | | X | *CTNNB1* | (39) |
| | | | | X | *CTNND2* | (39) |
| | | | | X | *CTTNBP2* | (39) |
| | | | | X | *CUL3* | (39) |
| X | | | | | *CUL3EP300* | (79) |
| | | | | X | *CUL7* | (39) |
| | | | | X | *CUX1* | (39) |
| | | | | X | *CYFIP1* | (39) |
| | | | | X | *CYP27A1* | (39), (64) |
| X | | | | | *DAO* | (83) |
| X | | X | | | *DAOA* | (41) |
| | | | | X | *DAPP1* | (39) |

| | | | | | | |
|---|---|---|---|---|---|---|
| | X | X | X | | *DAT1* | (56) |
| | | | X | | *DBH* | (56) |
| | | | | X | *DCC* | (39) |
| | X | | | | *DCLK3* | (49), (67) |
| | | | | X | *DDHD2* | (39) |
| | | | | X | *DDX3X* | (39) |
| | | | | X | *DEAF1* | (39) |
| | | | | X | *DENR* | (39) |
| | | | | X | *DEPDC5* | (39) |
| | X | | | | *DGKH* | (44) |
| | | | | X | *DHCR7* | (39) |
| | X | | | | *DHH* | (47), (67), (58) |
| | | | | X | *DHX30* | (39) |
| | | | | X | *DIP2A* | (39) |
| | | | | X | *DIP2C* | (39) |
| | | | X | | *DIRAS2* | (57) |
| X | | X | | X | *DISC1* | (39), (82) |
| | | | | X | *DLG2* | (39) |
| | | | | X | *DLG4* | (39) |
| | | | | X | *DLGAP1* | (39) |
| | | | | X | *Dlgap2* | (77) |
| | | | | X | *DLX3* | (39) |
| | | | | X | *DMD* | (39) |
| | | | | X | *DMPK* | (39) |
| | X | | | | *DNAH1* | (67), (49) |
| | | | | X | *DNMT3A* | (39) |
| | | | | X | *DOCK8* | (39) |
| | | | | X | *DOLK* | (39) |
| | | | X | | *DPH2* | (53) |
| | | | | X | *DPP10* | (39) |
| | | | | X | *Dpp6* | (66) |
| | | | | X | *DPYSL2* | (39) |
| | X | | | | *DRD1* | (73) |
| X | | | | | *DRD2* | (45) |
| | | | | X | *DRD3* | (80) |
| | X | X | X | | *DRD4* | (80) |
| | | | X | | *DRD5* | (70) |
| | | | | X | *DSCAM* | (39) |
| | | | X | | *DUSP6* | (53) |

| | | | | | | |
|---|---|---|---|---|---|---|
| | | | | X | *DYNC1H1* | (39) |
| | | | | X | *DYRK1A* | (39) |
| | | | | X | *EBF3* | (39) |
| | | | | X | *EEF1A2* | (39) |
| | | | | X | *EFR3A* | (39) |
| | | | | X | *EHMT1* | (39) |
| | | | | X | *EIF3G* | (39) |
| | | | | X | *ELAVL3* | (39) |
| | | | | X | *ELP4* | (39) |
| | | | | X | *EMSY* | (39) |
| | | | | X | *EP300* | (39) |
| | | | | X | *EP400* | (39) |
| X | | | | | *ERBB4* | (69) |
| | | | | X | *ERBIN* | (39) |
| X | | | | | *ESRP2* | (79) |
| | | | | X | *ETFB* | (39), (79) |
| | | | | X | *FAM92B* | (39) |
| | | | | X | *FBN1* | (39) |
| | | | | X | *FBRSL1* | (39) |
| X | | | | X | *FGFR1* | (39), (53) |
| | | | | X | *FMR1* | (39) |
| | | | | X | *FOXG1* | (39) |
| | | | | X | *FOXP1* | (39) |
| | | | X | X | *FOXP2* | (39) |
| | | | | X | *FRMPD4* | (39) |
| | X | | | | *FSTL5* | (81) |
| X | | | | | *FURIN* | (79) |
| | | | | X | *GABRA3* | (39) |
| | | | | X | *GABRB2* | (39) |
| | | | | X | *GABRB3* | (39) |
| | | | | X | *GABRG3* | (39) |
| | X | | | | *GAL3ST3* | (81) |
| | | | | X | *GALNT10* | (39) |
| | | | | X | *GALNT2* | (39) |
| | X | | | | *GALNT3* | (81) |
| | | | | X | *GALNT8* | (39) |
| | | | | X | *GATM* | (39) |
| | | | | X | *GFAP* | (39) |
| | | | | X | *GGNBP2* | (39) |

| | | | | | | |
|---|---|---|---|---|---|---|
| | | | | X | *GIGYF1* | (39) |
| | | | | X | *GIGYF2* | (39) |
| | | | X | | *GIT1* | (75) |
| | | | | X | *GNAI1* | (39) |
| X | | | | | *GNB3* | (65), |
| | | | | X | *GPC4* | (39) |
| | | | | X | *GPHN* | (39) |
| | X | | | | *GRB7* | (67), (58) |
| X | | | | X | *GRIA1* | (39), (43) |
| | | | | X | *Gria3* | (48) |
| | | | | X | *GRID1* | (39) |
| | | | | X | *GRIK2* | (39) |
| | | | | X | *GRIK5* | (39) |
| | | | | X | *GRIN1* | (39) |
| X | X | | | X | *GRIN2A* | (39) |
| | | | X | X | *GRIN2B* | (39) |
| | | | | X | *GRIP1* | (39) |
| | | | X | | *GRM1* | (57) |
| X | X | | | | *GRM3* | (60) |
| | | | | X | *HCFC1* | (39) |
| | | | | X | *HCN1* | (39) |
| | | | | X | *HDAC8* | (39) |
| | | | | X | *HDLBP* | (39) |
| | | | | X | *HECTD4* | (39) |
| | | | | X | *HECW2* | (39) |
| | | | | X | *HEPACAM* | (39) |
| | | | | X | *HERC2* | (39) |
| | | | | X | *HIVEP2* | (39) |
| | | | | X | *HIVEP3* | (39) |
| | | | | X | *HMGN1* | (39) |
| | | | | X | *HNRNPH2* | (39) |
| | | | | X | *HNRNPU* | (39) |
| | | | | X | *HOXA1* | (39) |
| | | | | X | *HRAS* | (39) |
| X | | | | | *HSPA9* | (79) |
| X | | | | | *HSPD1* | (79) |
| X | | | | | *HSPE1* | (79) |
| | | X | | | *HTR12A* | (80) |
| | | X | | | *HTR1A* | (80) |

| | | | | | | |
|---|---|---|---|---|---|---|
| | | X | | | *HTR1B* | (80) |
| | X | X | | | *HTR2A* | (65) |
| | | X | | | *HTR2C* | (80) |
| | X | | | | *HTTLPR* | (72) |
| | | | | X | *HUWE1* | (39) |
| | | | | X | *ICA1* | (39) |
| | | | | X | *ILF2* | (39) |
| X | | | | | *INA* | (79) |
| | | | | X | *INTS1* | (39) |
| | | | | X | *INTS6* | (39) |
| | | | X | | *IPO13* | (53) |
| | | | | X | *IQSEC2* | (39) |
| | | | | X | *IRF2BPL* | (39) |
| | | | | X | *ITGB3* | (39) |
| | | | | X | *JARID2* | (39) |
| | | | | X | *KANSL1* | (39) |
| | | | | X | *KAT2B* | (39) |
| X | | | | | *KAT5* | (79) |
| | | | | X | *KAT6A* | (39) |
| | | | | X | *KATNAL2* | (39) |
| | | | | X | *KCNB1* | (39) |
| | | | | X | *KCNJ10* | (39) |
| | | | | X | *KCNQ2* | (39) |
| | | | | X | *KCNQ3* | (39) |
| | | | | X | *KCNS3* | (39) |
| | | | | X | *KDM3B* | (39) |
| X | | | X | | *KDM4A* | (53) |
| | | | | X | *KDM4C* | (39) |
| | | | | X | *KDM5B* | (39) |
| | | | | X | *KDM5C* | (39) |
| | | | | X | *KDM6A* | (39) |
| | | | | X | *KDM6B* | (39) |
| | | | X | | *KDMA-AS1* | (53) |
| | | | | X | *KIAA0232* | (39) |
| | | | | X | *KIAA1586* | (39) |
| | | | | X | *KIF14* | (39) |
| | | | | X | *KIRREL3* | (39) |
| | | | | X | *KMT2A* | (39) |
| | | | | X | *KMT2C* | (39) |

| | | | | | | |
|---|---|---|---|---|---|---|
| | X | | | | *KMT2D* | (47), (67), (58) |
| | | | | X | *KMT2E* | (39) |
| | | | | X | *KMT5B* | (39) |
| | | | | X | *KPTN* | (39) |
| | | | | X | *LAMB1* | (39) |
| | | | | X | *LDB1* | (39) |
| | | | | X | *LEO1* | (39) |
| | | | X | | *LINC00461* | (53) |
| | X | | | | *LINC01215* | (81) |
| | | | X | | *LINC01288* | (53) |
| | | | X | | *LINC01572* | (53) |
| | | | X | | *LINC02060* | (53) |
| | | | X | | *LINC02497* | (53) |
| | | | | X | *LMX1B* | (39) |
| | | | | X | *LNPK* | (39) |
| | X | | | | *LOC101927314* | (71) |
| | X | | | | *LOC102725191* | (81) |
| | X | | | | *LOC105371789* | (81) |
| | | | X | | *LPHN3* | (79) |
| X | | | | X | *LRP1* | (39) |
| | | | | X | *LRRC4C* | (39) |
| X | | | | | *LSM1* | (79) |
| | | | | X | *LZTR1* | (39) |
| | | | | X | *MACROD2* | (39) |
| X | | | | | *MAD1L1* | (79) |
| | | | | X | *MAGEL2* | (39) |
| | | X | | | *MAOA* | (80) |
| | | | | X | *MAP1A* | (39) |
| | | | | X | *MAPT-AS1* | (39) |
| | | | | X | *MBD5* | (39) |
| | | | | X | *MBOAT7* | (39) |
| X | | | | | *MDK* | (79) |
| | | | | X | *MECP2* | (39) |
| | | | | X | *MED13* | (39) |
| | | | | X | *MED13L* | (39) |
| X | | | | X | *MEF2C* | (39) |
| | | | | X | *MEIS2* | (39) |
| | | | | X | *MET* | (39) |
| | | | | X | *MFRP* | (39) |

| | | | | | | |
|---|---|---|---|---|---|---|
| | | | | X | *MIEN1* | (67), (58) |
| X | | | | X | *MIR137* | (76) |
| | X | | | | *MIR3127* | (67), (49) |
| | | | X | | *MIR3666* | (53) |
| | X | | | | *MIR4655* | (67), (58) |
| | X | | | | *MIR4728* | (67), (58) |
| | X | | | | *MIR640* | (52), (67) |
| | | | X | | *MIR9-2* | (53) |
| | X | | | | *MIRLET7G* | (67), (49) |
| | | | | X | *MKX* | (39) |
| | | | | X | *MSL3* | (39) |
| | | | | X | *MSNP1AS* | (39) |
| X | | X | | | *MTHFR* | (65) |
| | | | | X | *MTOR* | (39) |
| | X | | | | *MXI1* | (47), (67) |
| | | | | X | *MYH10* | (39) |
| | | | | X | *MYH9* | (39) |
| | | | | X | *MYO5A* | (39) |
| | | | | X | *MYO9B* | (39) |
| | | | | X | *MYT1L* | (39) |
| | | | | X | *NAA15* | (39) |
| | | | | X | *NACC1* | (39) |
| | | | | X | *NAV2* | (39) |
| | | | | X | *NBEA* | (39) |
| | X | | | | *NCAM1* | (54) |
| X | X | | | | *NCAN* | (52), (67) |
| | | | | X | *NCKAP1* | (39) |
| | | | | X | *NCOA1* | (39) |
| | | | | X | *NCOR1* | (39) |
| X | | | | | *NDST3* | (64) |
| X | | | | | *NEK1* | (79) |
| | | | | X | *NEXMIF* | (39) |
| | | | | X | *NF1* | (39) |
| | | | | X | *NFE2L3* | (39) |
| | X | | | X | *NFIX* | (39) |
| X | | | | | *NGEF* | (79) |
| X | | | | | *NGN2/NEUROG2* | (51) |
| | | | | X | *NINL* | (39) |
| | | | | X | *NIPBL* | (39) |

| | | | | | | |
|---|---|---|---|---|---|---|
| | | | | X | *NLGN1* | (39) |
| | | | | X | *NLGN2* | (39) |
| | | | | X | *NLGN3* | (39) |
| | | | | X | *NLGN4X* | (39) |
| X | | | | | *NMB* | (79) |
| | | | X | | *NOS1* | (84) |
| X | | | | | *NOTCH4* | (64) |
| | | | | X | *NOVA2* | (39) |
| | | X | | | *NR3C1* | (65) |
| | | | | X | *NR3C2* | (39) |
| X | | X | | X | *NR4A2* | (39) |
| X | X | | | | *NRG1* | (68), (54) |
| X | | | | | *NRGN* | (79) |
| | | | | X | *NRXN1* | (39) |
| | | | | X | *NRXN2* | (39) |
| | | | | X | *NRXN3* | (39) |
| | | | | X | *NSD1* | (39) |
| | | | | X | *NSD2* | (39) |
| | | | | X | *NTNG2* | (39) |
| | | | | X | *NTRK2* | (39) |
| | | | | X | *NUAK1* | (39) |
| | | | | X | *NUDCD2* | (39) |
| | | | | X | *NUP155* | (39) |
| | | | | X | *OCRL* | (39) |
| | X | | | | *ODZ4* | (61) |
| X | | | | | *OGFOD2* | (79) |
| X | | | | | *OPCML* | (79) |
| | | | | X | *OPHN1* | (39) |
| | | | | X | *OR52M1* | (39) |
| | | | | X | *OTUD7A* | (39) |
| X | X | | | | *OTUD7B* | (79) |
| | | | | X | *OXTR* | (39) |
| | | | | X | *P2RX5* | (39) |
| | | | | X | *P4HA2* | (39) |
| | | | | X | *PACS1* | (39) |
| | | | | X | *PACS2* | (39) |
| | | | | X | *PAH* | (39) |
| | | | | X | *PAK1* | (39) |
| | | | | X | *PAK2* | (39) |

| | | | | | | |
|---|---|---|---|---|---|---|
| | | | | X | *PARD3B* | (39) |
| | | | X | | *PARK2* | (57) |
| | | | | X | *PAX5* | (39) |
| | | | | X | *PAX6* | (39) |
| | | | | X | *PCCA* | (39) |
| | | | | X | *PCCB* | (39) |
| | | | | X | *PCDH19* | (39) |
| | | | | X | *PCDH7* | (53) |
| | | | | X | *Pcdh9* | (66) |
| | | X | | | *PCLO* | (80) |
| | X | | | | *PDE10A* | (62) |
| X | | | | | *PDE4B* | (64) |
| | | | X | | *PDE4D* | (57) |
| | | | | X | *PER2* | (39) |
| X | | | | | *PGBD1* | (64) |
| X | | | | | *PGM3* | (79) |
| | | | | X | *PHB* | (39) |
| | | | | X | *PHF12* | (39) |
| | | | | X | *PHF2* | (39) |
| | | | | X | *PHF21A* | (39) |
| | | | | X | *PHF3* | (39) |
| | | | | X | *PHF7* | (39) |
| | | | | X | *PHF8* | (39) |
| | | | | X | *PHIP* | (39) |
| | | | | X | *PHRF1* | (39) |
| | | | | X | *PIK3R2* | (39) |
| | | | | X | *PLCB1* | (39) |
| | | | | X | *PLXNA4* | (39) |
| | | | | X | *PLXNB1* | (39) |
| | | | X | | *POC1B* | (53) |
| | | | | X | *POGZ* | (39) |
| | | | | X | *POMGNT1* | (39) |
| | | | | X | *PON1* | (39) |
| | | | | X | *POU3F3* | (39) |
| | | | | X | *PPP1R9B* | (39) |
| | | | | X | *PPP2CA* | (39) |
| X | | | | | *PPP2R2A* | (79) |
| | | | | X | *PPP2R5D* | (39) |
| | | | | X | *PPP5C* | (39) |

| | | | | | | |
|---|---|---|---|---|---|---|
| | | | | X | *PREX1* | (39) |
| | | | | X | *PRICKLE1* | (39) |
| | | | | X | *PRICKLE2* | (39) |
| | | | | X | *PRKCB* | (39) |
| | | | | X | *PRKD1* | (39) |
| | | | | X | *PRKD2* | (39) |
| | | | | X | *PRKN* | (39) |
| X | | | | | *PRMT1* | (79) |
| | | | | X | *PRODH* | (39) |
| | | | | X | *PRR12* | (39) |
| X | | | | | *PRSS16/POM121 L2* | (64) |
| | | | | X | *PSMD12* | (39) |
| | | | | X | *PTCHD1* | (39) |
| | | | | X | *PTCHD1-AS* | (39) |
| | | | | X | *PTEN* | (39) |
| X | | | | | *PTK2B* | (79) |
| | | | | X | *PTK7* | (39) |
| | | | | X | *PTPN11* | (39) |
| | | | X | | *PTPRF* | (53) |
| | | | | X | *PYHIN1* | (39) |
| | | | | X | *QRICH1* | (39) |
| | | | | X | *RAB2A* | (39) |
| | | | | X | *RAB43* | (39) |
| | | | | X | *RAC1* | (39) |
| | | | | X | *RAD21* | (39) |
| | | | X | X | *RAI1* | (39) |
| | | | | X | *RALA* | (39) |
| X | | | | | *RALGAPA2* | (79) |
| | | | | X | *RALGAPB* | (39) |
| | | | | X | *RANBP17* | (39) |
| X | | | | | *RANGAP1* | (79) |
| | | | | X | *RBFOX1* | (39) |
| | | | | X | *RBM27* | (39) |
| | X | | | | *RBPJL* | (81) |
| X | | | | | *RELA* | (79) |
| | | | | X | *RELN* | (39) |
| | | | | X | *RERE* | (39) |
| | | | | X | *RFX3* | (39) |
| | | | | X | *RHEB* | (39) |

| | | | | | | |
|---|---|---|---|---|---|---|
| | | | | X | *RHEBL1* | (47), (67), (58) |
| | | | | X | *RIMS1* | (39) |
| | | | | X | *RLIM* | (39) |
| | X | | | | *RNA5SP223* | (67) |
| | X | | | | *RNU2-17P* | (81) |
| | X | | | | *RNU6-1028P* | (52), (67) |
| | X | | | | *RNU6-131P* | (81) |
| | | | | X | *ROBO2* | (39) |
| | | | | X | *RORA* | (39) |
| | | | | X | *RORB* | (39) |
| | | | | X | *RPS10P2-AS1* | (39) |
| | X | | | | *RPS6KA2* | (62) |
| X | | | | | *RPTOR* | (79) |
| | | | | X | *RSRC1* | (39) |
| | | | | X | *SAE1* | (39) |
| | | | | X | *SATB1* | (39) |
| | | | | X | *SBF1* | (39) |
| | | | | X | *SCAF4* | (39) |
| | | | | X | *SCN1A* | (39) |
| | | | | X | *SCN2A* | (39) |
| | X | | | | *SCN2A* | (81) |
| | | | | X | *SCN8A* | (39) |
| | | | | X | *SCN9A* | (39) |
| | | | | X | *SEMA5A* | (39) |
| | | | X | | *SEMA6D* | (53) |
| X | | | | | *SERPING1* | (79) |
| | | | | X | *SET* | (39) |
| | | | | X | *SETBP1* | (39) |
| | | | | X | *SETD1A* | (39) |
| | | | | X | *SETD2* | (39) |
| | | | | X | *SETD5* | (39) |
| X | | | | | *SF3B1* | (79) |
| | | | | X | *SGSH* | (39) |
| | | | | X | *SHANK1* | (39) |
| | X | | | X | *SHANK2* | (39) |
| | | | | X | *SHANK3* | (39) |
| | | | | X | *SHOX* | (39) |
| | | | | X | *SIK1* | (39) |
| | | | | X | *SIN3A* | (39) |

| | | | | | | |
|---|---|---|---|---|---|---|
| | | | | X | *SKI* | (39) |
| | | | | X | *SLC12A5* | (39) |
| | | | | X | *SLC1A2* | (39) |
| | | | | X | *SLC35B1* | (39) |
| | | | | X | *SLC38A10* | (39) |
| | | | | X | *SLC45A1* | (39) |
| | | | | X | *SLC6A1* | (39) |
| | | X | | | *SLC6A2* | (65) |
| | | X | X | X | *SLC6A3* | (39) |
| | X | X | X | | *SLC6A4* | (54) |
| | | | X | | *SLC6A9* | (53) |
| | | | | X | *SLC7A3* | (39) |
| | | | | X | *SLC7A5* | (39) |
| | | | | X | *SLC9A6* | (39) |
| | | | | X | *SLITRK5* | (39) |
| | | | | X | *SMAD4* | (39) |
| | | | | X | *SMARCA2* | (39) |
| | | | | X | *SMARCA4* | (39) |
| | | | | X | *SMARCC2* | (39) |
| | | | | X | *SMC1A* | (39) |
| | X | | | | *SMNDC1* | (47), (67) |
| | | | X | | *SNAP25* | (70) |
| X | | | | | *SNAP91* | (51) |
| | | | | X | *SNX14* | (39) |
| | | | | X | *SNX5* | (39) |
| | | | | X | *SON* | (39) |
| | X | | | | *SORCS1* | (47), (67) |
| | | | X | X | *SORCS3* | (39) |
| | | | | X | *SOX5* | (39) |
| | | | | X | *SOX6* | (39) |
| | | | X | | *SPAG16* | (53) |
| | | | | X | *SPARCL1* | (39) |
| | | | | X | *SPAST* | (39) |
| | | | | X | *SPEN* | (39) |
| | | | | X | *SRCAP* | (39) |
| X | | | | | *SREBF1* | (79) |
| X | | | | | *SRPK2* | (79) |
| | | | | X | *SRPRA* | (39) |
| X | | | | | *SRR* | (42) |

| | | | | | | |
|---|---|---|---|---|---|---|
| | | | | X | *SRSF11* | (39) |
| | | | X | | *ST3GAL3* | (53) |
| | | | | X | *ST8SIA2* | (39) |
| | | | | X | *STAG1* | (39) |
| | | | | X | *STXBP1* | (39) |
| | | | | X | *STXBP5* | (39) |
| | | | | X | *SUPT16H* | (39) |
| | | | | X | *SYN1* | (39) |
| X | | | | | *Syn3* | (50) |
| | X | | | | *SYNE1* | (67) |
| | X | | | | *SYNE1-AS1* | (67) |
| | | | | X | *SYNGAP1* | (39) |
| | | | | X | *SYT1* | (39) |
| | | | X | | *SYT2* | (57) |
| | | | | X | *TAF1* | (39) |
| | | | | X | *TAF6* | (39) |
| | | | | X | *TANC2* | (39) |
| | | | | X | *TAOK1* | (39) |
| | | | | X | *TAOK2* | (39) |
| | | | | X | *TBC1D23* | (39) |
| | | | | X | *TBC1D31* | (39) |
| | | | | X | *TBCK* | (39) |
| | | | | X | *TBL1XR1* | (39) |
| | | | | X | *TBR1* | (39) |
| | | | | X | *TBX1* | (39) |
| | | | | X | *TCF20* | (39) |
| X | | | | X | *TCF4* | (39) |
| | | | | X | *TCF7L2* | (39) |
| | | | | X | *TEK* | (39) |
| | | | | X | *TERF2* | (39) |
| | | | | X | *TET2* | (39) |
| | | | | X | *TET3* | (39) |
| | | X | | | *TH* | (80) |
| | X | | | | *THSD7A* | (81) |
| | | | | X | *TLK2* | (39) |
| | X | | | | *TLR9* | (49), (67) |
| | | | | X | *TM4SF20* | (39) |
| | | | | X | *TM9SF4* | (39) |
| | X | | | | *TMEM151A* | (81) |

| | | | | | | |
|---|---|---|---|---|---|---|
| | | | | X | *TMLHE* | (39) |
| | X | | | | *TMUB2* | (81) |
| | | | | X | *TNRC6B* | (39) |
| | | | | X | *tor3a* | (63) |
| | | X | X | | *TPH1* | (75) |
| | | X | X | | *TPH2* | (75) |
| | | | | X | *TRAF7* | (39) |
| | X | | | | *TRANK1* | (49), (67) |
| | | | | X | *TRAPPC6B* | (39) |
| | | | | X | *TRAPPC9* | (39) |
| | | | | X | *TRIM23* | (39) |
| | | | | X | *TRIO* | (39) |
| | | | | X | *TRIP12* | (39) |
| | | | | X | *TRPC6* | (39) |
| | | | | X | *TRPM1* | (39) |
| | | | | X | *TSC1* | (39) |
| | | | | X | *TSC2* | (39) |
| | | | | X | *TSHZ3* | (39) |
| X | | | | | *TSNARE1* | (51) |
| | | | | X | *TTI2* | (39) |
| | | | | X | *UBE3A* | (39) |
| | | | | X | *UBE3C* | (39) |
| | | | | X | *UBN2* | (39) |
| | | | | X | *UBR1* | (39) |
| | | | | X | *UBR5* | (39) |
| | | | | X | *UNC79* | (39) |
| | | | | X | *UPF3B* | (39) |
| | | | | X | *USP15* | (39) |
| | | | | X | *USP45* | (39) |
| | | | | X | *USP7* | (39) |
| | | | | X | *USP9X* | (39) |
| | | | | X | *VAMP2* | (39) |
| | | | | X | *VEZF1* | (39) |
| | | | | X | *VIL1* | (39) |
| | | | | X | *VPS13B* | (39) |
| | | | | X | *WAC* | (39) |
| | | | | X | *WASF1* | (39) |
| | | | | X | *WDFY3* | (39) |
| | | | | X | *WDFY4* | (39) |

| | | | | | | |
|---|---|---|---|---|---|---|
| | | | | X | *WDR26* | (39) |
| | | | | X | *WFDC5* | (81) |
| | | | | X | *WWOX* | (39) |
| | | | | X | *XPC* | (39) |
| | | | | X | *YY1* | (39) |
| | | | | X | *ZBTB20* | (39) |
| | | | | X | *ZC3H4* | (39) |
| | X | | | | *ZCCHC2* | (81) |
| X | | | | | *ZEB2* | (79) |
| | | | | X | *ZMIZ1* | (39) |
| | | | | X | *ZMYM2* | (39) |
| | | | | X | *ZMYND11* | (39) |
| | | | | X | *ZMYND8* | (39) |
| | | | | X | *ZNF292* | (39) |
| | | | | X | *ZNF462* | (39) |
| X | | | | X | *ZNF804A* | (39) |
| | | | | X | *ZSWIM6* | (39) |
